# Growth hormone is required for hippocampal engram cell maturation

**DOI:** 10.1101/2025.02.27.640537

**Authors:** Chang-Ho Kim, HyoJin Park, Dae Hee Han, Ilgang Hong, Yeonjun Kim, Nam-Kyung Yu, Jun Cho, Bong-Kiun Kaang

## Abstract

Memory is thought to be stored in a sparse population of neurons or synapses ^1^. These neurons or synapses collectively termed the engram are necessary and sufficient for memory recall ^2,3^. Learning induces synaptic strengthening in engram cells at both pre- and postsynaptic sites ^4,5^. However, the critical time window and the molecule related to the induction of such synaptic changes are unknown. Here we show that the initial translation plays a key role in engram maturation by facilitating pre- and postsynaptic strengthening and that growth hormone (GH) is required to induce this process in the Dentate Gyrus. Using anisomycin, a protein synthesis inhibitor, we found that blocking the initial and the subsequent second wave of translation arrests the maturation of engram cells. Revisiting our hippocampal translatome during memory formation ^6^ proposed GH as a mediator of engram maturation. Overexpression of the dominant negative (G118R) GH blocked the maturation of the hippocampal engram cell ^7^. Facilitating activity-dependent GH uptake by injecting recombinant human GH (rhGH) into the animal rescued the arrestment caused by anisomycin. Together, our findings propose GH as a key mediator of hippocampal engram cell maturation.

## 1. Main

The engram represents the physical substrate of memory ^1^. Previous studies have identified that the activity of engram cells is necessary and sufficient for memory recall ^2,3^. We and others have questioned the state of the engram before memory formation and found that the properties (synaptic connectivity and neuronal excitability) of the engram cell are different from those of non-engram cells ^8-10^. This suggests that a distinct subset of cells, characterized by unique properties, undergoes maturation in a learning-dependent manner, thereby strengthening its functional attributes and contributing to the formation of the engram. Nevertheless, the critical time window and the molecular pathways required for the maturation of engram cells remain elusive.

De novo protein synthesis is a critical mechanism of long-term memory formation, represented by impaired retrieval of long-term memory once the protein synthesis is blocked at the time of learning ^4,11-17^. However, despite the protein synthesis blockade, several properties of the engram remain intact. The sufficiency of the hippocampal engram cell for memory recall was not disturbed under short inhibition of the translational process ^4^. By extending the inhibition, we previously found that the sufficiency of circuit-level activation in the ventral CA1 to the basal amygdala was impaired ^14^. However, in both behavioral paradigms, engram-specific presynaptic strengthening and the increase in the density of engram synapses were observed, indicating functional maturation of engram cells ^4,14,18^.

To identify the time window for the maturation of engram cells, we revisited the nature of learning. Intuitively, learning does not occur after the termination of a specific experience. It begins during the experience. Such characteristic is represented by differences in memory depending on the timing of protein synthesis inhibition ^11^. Given that the properties of the engram were partially intact when protein synthesis was inhibited after learning ^4,14,18-20^, we hypothesized that the critical translational processes for engram maturation might occur immediately at the time of learning. To test the hypothesis, we divided memory consolidation into three distinct phases: the initial phase (corresponding to the learning period), the early phase (immediately following the learning period), and the late phase (associated with the second wave of translation). We then disrupted protein synthesis in these phases by administrating anisomycin intraperitoneally (i.p.) at various time points relative to the contextual fear conditioning (CFC) process. In the [Multiple (−1 hr)] group, all three phases were disrupted, whereas in the other groups, no phases [Saline], a single phase [Single (0 hr)], or two phases [Single (−1 hr) and Multiple (0 hr)] were selectively disrupted (Fig. 1a and extended data fig. 1a, 3a). We targeted the Dentate Gyrus (DG) as our brain region of interest as the engram cells in the DG were sufficient for memory recall under protein synthesis blockade ^4^. To identify the changes in the dendritic spine density, engram cells were labeled in an activity (c-fos) and doxycycline-dependent (rtTA3G and TRE3G) manner. Non-engram cells were identified among the randomly and sparsely labeled cells using the Cre recombinase virus that were not labeled by activity (Fig. 1b and extended data fig. 1b, 2a-b) ^5^. The effect of anisomycin was minimal on the virally expressed protein as the difference between the groups disappeared by 4 hours following the CFC (extended data fig. 2d-g). Furthermore, 2 days after labeling, the number of labeled cells was indistinguishable among the groups (extended data fig. 2h-i). In line with previous studies, long-term memory tested 2 days after the CFC was impaired under all conditions where anisomycin was administered (Fig. 1c and extended data fig. 1c) ^4,14,20^. Notably, the freezing response in the Multiple (−1 hr) group was indistinguishable between the pre-exposure and the retrieval session. Analysis of the dendritic spine density indicated that engram dendrites in the Multiple (−1 hr) group had lower spine density than non-engram dendrites (Fig. 1d-e and extended data fig. 1d-e). This difference was not due to artifacts caused by fluorescence proteins (extended data fig. 2j-k). These results are reminiscent of our previous study, in which cells that eventually became the engram had less synaptic connection than non-engram cells before learning ^10^.

**Figure. 1:**
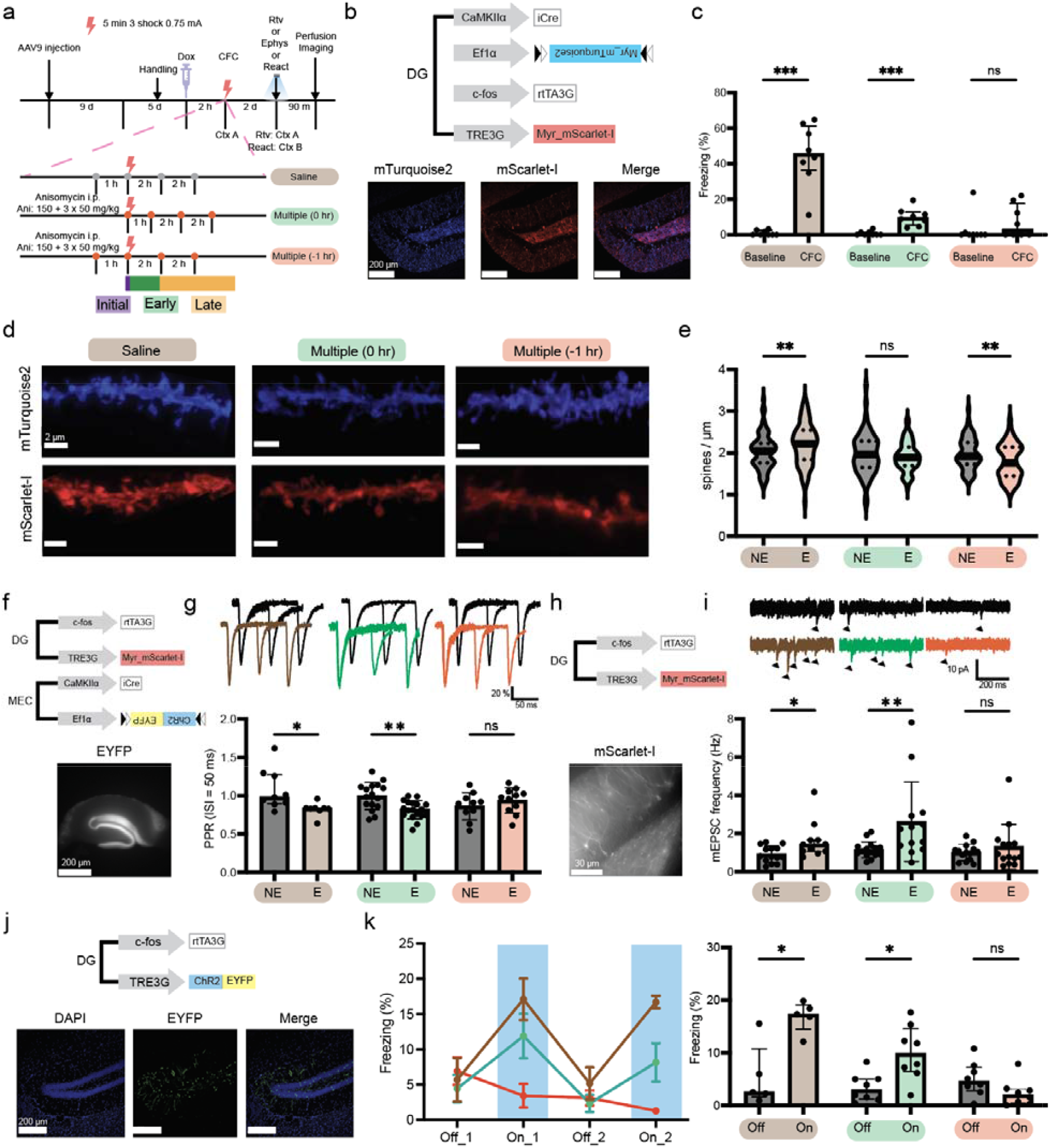
Blocking the protein synthesis during the acquisition and the subsequent second wave of translation after learning arrests the maturation of hippocampal engram cells. **a**, Experimental procedures for the behavioral paradigm used to arrest engram cell maturation. **b**, Top: viral constructs for natural memory retrieval and spine density analysis. Bottom: image of the fluorescent proteins in the DG. **c**, Results of the natural memory retrieval (n = 8 animals per group). **d**, Image of the dendrite segment. **e**, Dendritic spine density of neurons (from left to right, n = 83, 84, 72, 73, 64, 68 dendrites). **f**, Top: viral constructs for PPR analysis. Bottom: Image of the labeled medial perforant pathway. **g**, Top: traces of the PPR. Bottom: PPR (ISI = 50 ms) of neurons (from left to right, n = 9, 8, 16, 18, 10, 11 neurons). **h**, Top: viral constructs for mEPSC analysis. Bottom: image of engram neurons. **i**, Top: traces of the mEPSC. Black arrows indicate each mEPSC spike. Bottom: mEPSC frequency of neurons (from left to right, n = 11, 14, 15, 13, 14, 14 neurons). **j**, Top: viral constructs for artificial engram reactivation. Bottom: image of the DG expressing the fluorescent protein. **k**, Left: behavior paradigm and the quantified freezing behavior. Right: the quantified freezing behavior (from left to right, n = 5, 8, 8 animals). Mann Whitney U test, ***: P < 0.001, **: P < 0.01, *: P < 0.05, ns: P > 0.05.

**Figure. 2:**
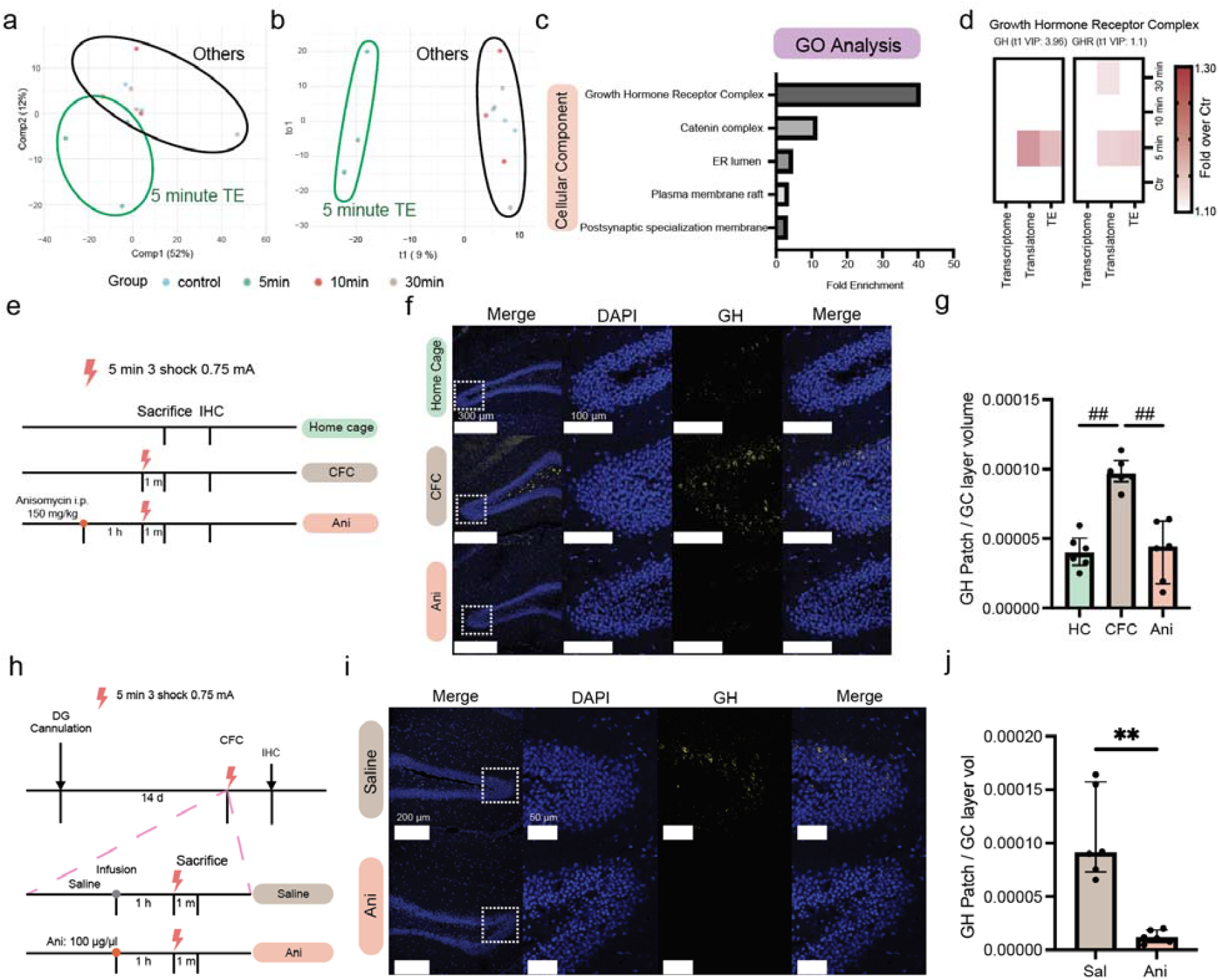
Growth hormone is rapidly translated in the hippocampus during learning. **a**, Low dimensional analysis of the TE using Multilevel-PCA. **b**, OPLS-DA on the TE. **c**, Fold enrichment of significantly altered GO terms (FDR < 0.05). **d**, Heatmap of genes enriched in the growth hormone receptor complex. **e**, Experimental procedures for the GH-IHC. **f**, Images of the GH-IHC. **g**, Quantified patches of GH in the granule cell layer (n = 6 animals per group). **h**, Experimental procedures for the GH-IHC. **i**, Images of the GH-IHC. **j**, Quantified patches of GH in the granule cell layer (n = 6 animals per group). Mann Whitney U test, **: P < 0.01. Dunn test, ##: adj.P < 0.01.

Next, we tested whether the initial protein synthesis has a key role in the presynaptic properties of the engram cells. The synaptic input from the medial entorhinal cortex (MEC) to the DG (Medial Perforant Pathway, MPP) is known to transmit contextual information associated with an experience ^21,22^. Therefore, we labeled the excitatory connections from the MEC to the DG using the channelrhodopsin-2 (ChR2) and measured the paired-pulse ratio (PPR) as a proxy for the presynaptic release probability (Fig. 1f and extended data fig. 1f). In all experimental conditions other than the Multiple (−1 hr) group, engram cells showed a reduced PPR at the 50 ms inter-stimulus-interval (ISI), but not at the 100 ms ISI suggesting presynaptic strengthening occurred through learning (Fig. 1g and extended data fig. 1g, 3b). However, engram-specific synaptic plasticity was not observed in the Multiple (−1 hr) group. To confirm the changes in release probability, we examined the difference in miniature excitatory postsynaptic current (mEPSC) frequency (Fig. 1h and extended data fig. 1h) ^23^. In accordance with the PPR data, engram-specific increase in mEPSC frequency was observed in all groups except the Multiple (−1 hr) group (Fig. 1i and extended data fig. 1i).

To assess differences in the postsynaptic properties, we measured the AMPA/NMDA ratio (MPP stimulation) and the capacitance of engram and non-engram cells. Consistent with a previous study, anisomycin administration resulted in indistinguishable properties between engram and non-engram cells, regardless of the disrupted consolidation phase (extended data fig. 3c-d) ^4^. These results were not due to artifacts caused by viral overexpression or doxycycline injections (extended data fig. 2c).

**Figure. 3:**
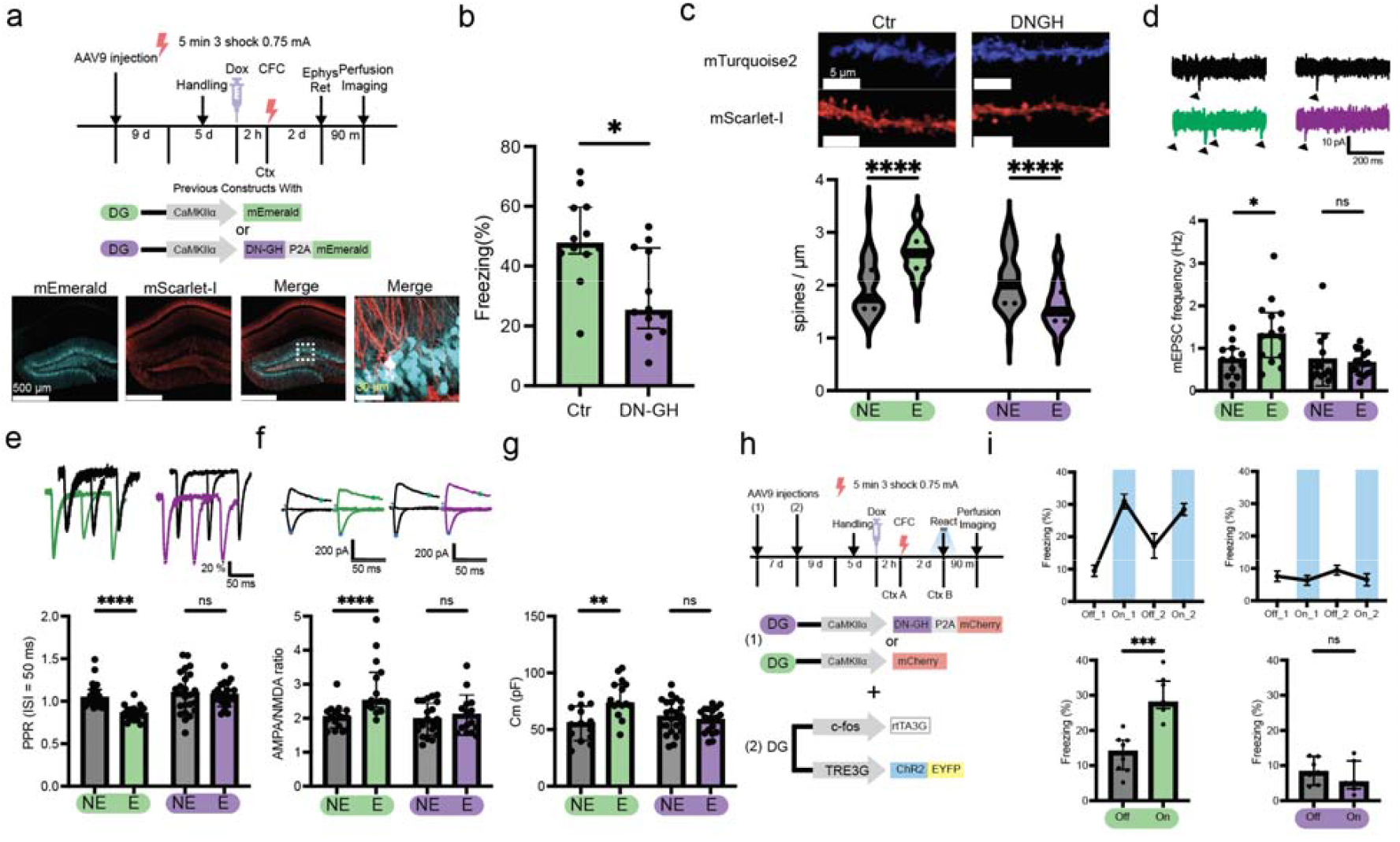
Growth hormone is required for the maturation of engram cells. **a**, Top: Experimental procedures for the behavioral paradigm. Middle: viral constructs used. Bottom: image of the DG expressing mScarlet-I and DN-GH-P2A-mEmerald. **b**, Results of the natural memory retrieval (n = 12 animals per group). **c**, Top: image of the dendrite segment. Bottom: dendritic spine density of neurons (from left to right, n = 41, 41, 51, 50 dendrites). **d**, Top: traces of the mEPSC. Black arrows indicate each mEPSC spike. Bottom: mEPSC frequency of neurons (from left to right, n = 12, 14, 13, 16 neurons). **e**, Top: traces of the PPR. Bottom: PPR (ISI = 50 ms) of neurons (from left to right, n = 23, 17, 24, 22 neurons). **f**, Top: AMPA/NMDA ratio traces. Bottom: AMPA/NMDA ratio of neurons (from left to right, n = 17, 15, 21, 15). **g**, Capacitance of neurons (from left to right, n = 14, 15, 23, 19 neurons). **h**, Top: Experimental procedures for the behavioral paradigm. Bottom: viral constructs used. **i**, Top: behavior paradigm and the quantified freezing behavior. Bottom: the quantified freezing behavior (Ctr: n = 8, DN-GH: n = 7). Mann Whitney U test, ****: P < 0.0001, ***: P < 0.001, **: P < 0.01, *: P < 0.05, ns: P > 0.05. Dunn test, ##: adj.P < 0.01.

We speculated that the maturation of engram cells indicated by engram-specific synaptic plasticity would be crucial for the sufficiency of engram for memory recall. As shown in previous studies, optogenetic activation of the engram cells using the ChR2 elicited aversive memory-dependent freezing behaviors (extended data fig. 4a-b) ^2,4^. Then, we confirmed that anisomycin injections do not significantly alter the functionality of ChR2 through c-fos immunohistochemistry (IHC) and ex vivo electrophysiology (extended data fig. 4c-i). Lastly, we tested whether the sufficiency of the engram for memory recall was altered (Fig. 1j and extended data fig. 1j). As expected, all behavioral groups except for the Multiple (−1 hr) groups showed significantly higher freezing responses on the light On sessions than the Off sessions (Fig. 1k and extended data fig. 1k).

**Figure. 4:**
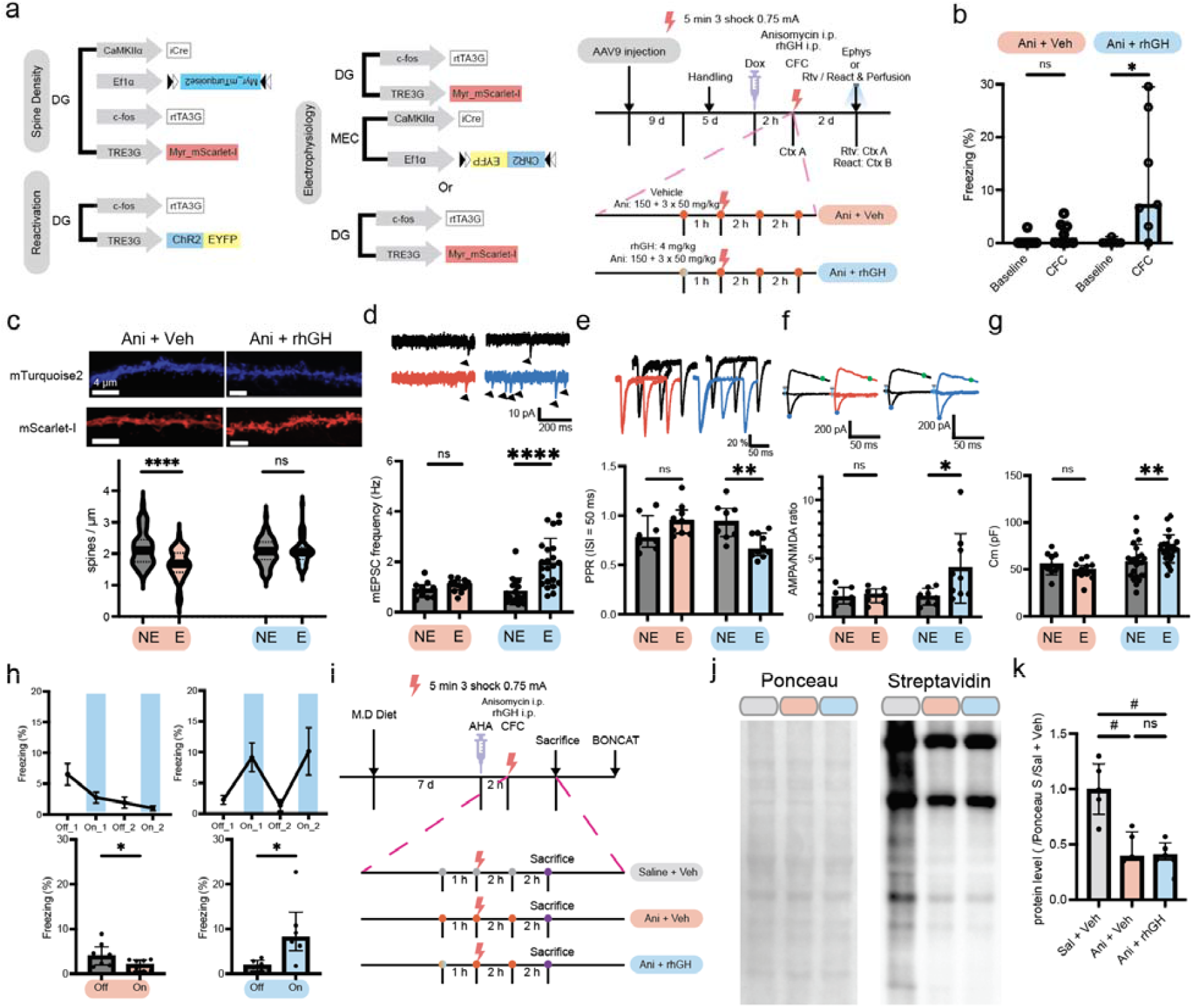
Growth hormone rescues anisomycin-induced arrestment of engram cell maturation and memory. **a**, Left: viral constructs used. Right: experimental procedures for the rhGH-mediated rescue. **b**, Results of the natural memory retrieval (Ani+veh: n = 8, Ani+rhGH: n = 7 animals). **c**, Top: image of the dendrite segment. Bottom: dendritic spine density of neurons (from left to right, n = 61, 60, 67, 71 dendrites). **d**, Top: traces of the mEPSC. Black arrows indicate each mEPSC spike. Bottom: mEPSC frequency of neurons (from left to right, n = 10, 12, 17, 23 neurons). **e**, Top: traces of the PPR. Bottom: PPR (ISI = 50 ms) of neurons (from left to right, n = 8, 9, 8, 10 neurons). **f**, Top: AMPA/NMDA ratio traces. Bottom: AMPA/NMDA ratio of neurons (from left to right, n = 7, 7, 8, 9). **g**, Capacitance of neurons (from left to right, n = 11, 12, 24, 24 neurons). **h**, Top: behavior paradigm and the quantified freezing behavior. Bottom: the quantified freezing behavior (Ani+veh: n = 8, Ani+rhGH: n = 7 animals). **i**. Experimental procedures for BONCAT. **j**, Image of the western blot. **k**, Quantified protein synthesis (n = 5 animals per group). Mann Whitney U test, ****: P < 0.0001, **: P < 0.01, *: P < 0.05, ns: P > 0.05. Dunn test, #: adj.P < 0.05, ns: adj.P > 0.05.

Collectively, in the Multiple (−1 hr) group, the properties of the mature engram were not observed. This suggests that blocking all three consolidation phases (initial, early, and late) arrests the maturation of hippocampal engram cells.

Next, we explored the fundamental molecular mechanisms that drive cellular maturation into engrams. Since the results from anisomycin injections indicated that the key translational regulation occurs during or immediately after learning, we reasoned that the regulation may occur through changes in translational efficiency (TE). Thus, we re-analyzed our data on the mouse hippocampus’s learning-associated changes in TEs ^6^. Multilevel-principal component analysis (multilevel-PCA^24^) on the genome-wide TE data revealed distinct changes in TEs at the 5 min following learning (Fig. 2a). Subsequently, we performed orthogonal-partial least squared analysis (OPLS-DA) on the TE data (5 min vs others) and acquired the t1 (orthogonal component) variable influence in projection (VIP) score (Fig. 2b). Finally, we classified the genes based on the VIP score (VIP > 1.1) and fold change (5 min/others > 1.1), and conducted functional annotation analysis (https://geneontology.org/) ^25^. The results overrepresented the GH receptor complex, proposing the complex as the critical mediator of engram cell maturation (Fig. 2c). Given that GH harbored the highest VIP score within the complex (Fig. 2d), it was selected as the primary focus of our analysis. This overrepresentation appeared only in the translatome while not in the transcriptome (Fig. 2d), suggesting that this regulation occurs at the level of translation.

GH is known to be up-regulated after learning, modulate memory, cause postsynaptic cLTP, increase mEPSC frequency and spine density, and allocate engram cells ^26-31^. To address whether GH may play an important role in engram maturation, we asked whether the hippocampal GH is translated in a learning-dependent manner. We sacrificed animals immediately after the CFC and performed GH-IHC (Fig. 2e). We noticed translation- and CFC-dependent rise of sparse cell body-like patches of GH (Fig. 2f-g), which was similar to the sparse nature of engram cells in the DG (extended data fig. 2h-i) ^2^. Subsequently, we asked if the GH is translated in the DG through drug infusion experiments (Fig. 2h). Infusing anisomycin directly into the DG blocked the rise in GH signals (Fig. 2i-j). These findings indicate that GH is rapidly translated in the DG during the learning process.

To test if the GH is required for the maturation of engram cells, we disrupted the GH-mediated signaling by overexpressing the dominant negative (G118R) GH (DN-GH) in the DG (Fig. 3a) ^7^. The overexpression significantly impaired contextual memory and disrupted the synaptic changes of the engram as indicated by the dendritic spine density, and electrophysiological properties (mEPSC frequency, PPR, AMPA/NMDA ratio, and capacitance) (Fig. 3b-g). These changes were specific to the engram cell, and the number of labeled cells was not altered (extended data fig.5). Furthermore, we found that the sufficiency of the engram for memory recall was disrupted by the DN-GH (Fig. 3h-i). Expressing DN-GH in the MEC did not alter the PPR changes of the engram cell (extended data fig. 6). Collectively, disrupting postsynaptic GH-mediated signaling results in a similar phenotype with the Multiple (−1 hr) anisomycin injections, highlighting its requirement for the maturation of engram cells.

Notably, intracellular GH function was required for the presynaptic changes (PPR and mEPSC frequency) of engram cells, the properties identified as the major difference between the Multiple (−1 hr) and Multiple (0 hr) groups. However, previous studies have shown that while rhGH in bath ACSF does not affect the PPR, its i.p. administration induces alteration in both PPR and mEPSC frequency ^28,29,32^. One potential explanation is that intracellular GH exerts distinct functional effects, and when external GH is introduced via i.p. injection, it may be internalized to mediate these effects. We further hypothesized that this process parallels that of brain-derived neurotrophic factors (BDNFs), which are internalized in response to neuronal activity ^33^. Supporting this notion, during the early postnatal period (P14-21), when hippocampal inhibitory circuits remain immature and engram cells are not sparse, i.p. administration of rhGH resulted in a global increase in mEPSC frequency ^29,34^. To test if internalization of external GH occurred, we incubated the brain slices in His-tag rhGH for 10 minutes and stimulated the MPP with four rounds of intermittent theta burst stimulation (4X iTBS). Subsequently, we performed His tag-IHC, and the results showed that the His-tag is highly localized in the stimulated granule cell layer (extended data fig. 7a). Furthermore, we examined whether the internalized rhGH caused presynaptic strengthening (extended data fig. 7b). Notably, cells stimulated with iTBS under bath rhGH showed lower PPR than those stimulated without rhGH or those incubated with rhGH but without stimulation (extended data fig. 7c). In line with our PPR data, the sEPSC frequency was significantly higher for those stimulated under bath rhGH (extended data fig. 7d).

Engram cells are highly active during memory formation. If the activity of the engram cell is sufficient for external GH internalization, acute administration of rhGH would potentially alleviate the maturation arrestment caused by anisomycin. To examine this scenario, we tested whether rhGH administration reverses the inhibitory effect of anisomycin on engram cell maturation (Fig. 4a). Those injected with rhGH showed significantly higher freezing levels than baseline. (Fig. 4b). The dendritic spine density of engram cells was partially recovered by the rhGH injection (Fig. 4c). Furthermore, the rhGH injection rescued the electrophysiological properties blocked by anisomycin (mEPSC frequency, PPR, AMPA/NMDA ratio, and capacitance) (Fig. 4d-g). Lastly, the sufficiency of the engram cell for memory recall was also restored (Fig. 4h). The changes in the spine density and electrophysiological properties were specific to the engram, and the number of labeled cells was not altered by the rhGH injections (extended data fig. 8). Using biorthogonal noncanonical amino acid tagging (BONCAT), we confirmed that the effect of GH was not due to the disinhibition of the general translational process in the hippocampus (Fig.4i-k) ^35-37^. These results demonstrate that rhGH rescues anisomycin-mediated arrest of engram cell maturation.

## 2. Discussion

The sufficiency for memory recall is a defining trait of the engram ^2^. By extending (6 + hours) the protein synthesis block, we have previously identified the synaptic engram as a neural correlate of memory recall at the circuit level via axon terminal activation ^14^. However, the cellular correlate of memory recall through cellular activation and the mechanism for engram cell maturation remained elusive ^4,20^. This study demonstrated that the key translational mechanism driving engram cell maturation occurs immediately during the learning process, marked by significant alterations in spine density, capacitance, mEPSC frequency, AMPA/NMDA ratio, and PPR once it is blocked. Next, we have shown that GH is synthesized in a learning-dependent manner and is required to develop these properties of engram cells. Lastly, we showed that rhGH can be internalized through neural activity, which can rescue the anisomycin-induced maturation arrestment. Collectively, this study highlights GH as a critical mediator of engram cell maturation.

GH is synthesized in both the pituitary gland and the hippocampus ^30,38-40^. While neurons appear capable of internalizing externally derived GH (extended data fig. 7), evidence suggests that the hippocampus serves as the principal source of GH required for hippocampal engram maturation under physiological conditions. Previous studies have identified GH mRNA transcription as being regulated by the cytoplasmic polyadenylation element-binding protein 1 (CPEB-1), with a rapid increase in hippocampal GH mRNA observed immediately following learning ^30,39^. Supporting this, intrahippocampal anisomycin infusion blocked the upregulation of GH patches within the DG (Fig. 2h-j), further indicating that GH primarily originates locally within the hippocampus.

## Methods

### Animals

All experiments used male C57BL/6J mice purchased from DBL Korea. Female mice were not used due to the effects of estrogen levels on the growth hormone ^40^. The age of the animals at the time of the stereotaxic surgery was 3-4 weeks for electrophysiology and 6-8 weeks for other experiments (spine density, natural retrieval, and optogenetic engram reactivation). Mice were given ad libitum access to food and water and were raised in a 12-hr light/dark cycle. Mice were caged together (maximum of 5) until the day before handling and were single-caged afterward. An enriched environment was not provided for the full duration of the experiments. All procedures and animal care followed the regulations and guidelines of the Institutional Animal Care and Use Committees (IACUC) of the Institute for Basic Science.

### Virus-mediated gene expression

The recombinant AAV vectors were packaged with AAV9 coat proteins in the IBS Virus Facility (https://centers.ibs.re.kr/html/virusfacility_en/). For spine density analysis, the titers were 1.6 × 10^12^ genome copy (GC) mL^-1^ for AAV9-TRE3G-myr-mScarlet-I, 6 × 10^11^ GC mL^-1^ for AAV9-c-fos-rtTA3G, 1 × 10^9^ GC mL^-1^ for AAV9-CamKIIα-iCre, and 1.2 × 10^12^ GC mL^-1^ for AAV9-EF1α-DIO-myr-mTurquois2. For mEPSC analysis, the titers were 1.6 × 10^12^ GC mL^-1^ for AAV9-TRE3G-myr-mScarlet-I and 6 × 10^11^ GC mL^-1^ for AAV9-c-fos-rtTA3G. For PPR and AMPA/NMDA ratio analysis, the titers were 1.6 × 10^12^ GC mL^-1^ for AAV9-TRE3G-myr-mScarlet-I, 6 × 10^11^ GC mL^-1^ for AAV9-c-fos-rtTA3G, 6.25 × 10^11^ GC mL^-1^ for AAV9-CamKIIα-iCre, and 1 × 10^12^ GC mL^-1^ for AAV9-EF1α-DIO-ChR2-EYFP. For viral overexpression control experiments, the titers were 1.6 × 10^12^ GC mL^-1^ for AAV9-TRE3G-myr-mScarlet-I and 6 × 10^11^ GC mL^-1^ for AAV9-CamKIIα -rtTA3G. For general optogenetic engram reactivation experiments, the titers were 7.5 × 10^12^ GC mL^-1^ for AAV9-TRE3G-ChR2-EYFP and 7.5 × 10^12^ GC mL^-1^ for AAV9-c-fos-rtTA3G. For DN-GH mEPSC analysis, the titers were 1.6 × 10^12^ GC mL^-1^ for AAV9-TRE3G-myr-mScarlet-I, 4 × 10^12^ GC mL^-1^ for AAV9-CamKIIα-DN-GH-P2A-mEmerald, and 6 × 10^11^ GC mL^-1^ for AAV9-c-fos-rtTA3G. For control experiments, the titers were 1.6 × 10^12^ GC mL^-1^ for AAV9-TRE3G-myr-mScarlet-I, 4 × 10^12^ GC mL^-1^ for AAV9-CamKIIα-mEmerald, and 6 × 10^11^ GC mL^-1^ for AAV9-c-fos-rtTA3G. For DN-GH PPR and AMPA/NMDA ratio analysis, the titers were 1.6 × 10^12^ GC mL^-1^ for AAV9-TRE3G-myr-mScarlet-I, 6 × 10^11^ GC mL^-1^ for AAV9-c-fos-rtTA3G, 4 × 10^12^ GC mL^-1^ for AAV9-CamKIIα-DN-GH-P2A-mEmerald, 6.25 × 10^11^ GC mL^-1^ for AAV9-CamKIIα-iCre, and 1 × 10^12^ GC mL^-1^ for AAV9-EF1α-DIO-ChR2-EYFP. For control experiments, the titers were 1.6 × 10^12^ GC mL^-1^ for AAV9-TRE3G-myr-mScarlet-I, 6 × 10^11^ GC mL^-1^ for AAV9-c-fos-rtTA3G, 4 × 10^12^ GC mL^-1^ for AAV9-CamKIIα-mEmerald, 6.25 × 10^11^ GC mL^-1^ for AAV9-CamKIIα-iCre, and 1 × 10^12^ GC mL^-1^ for AAV9-EF1α-DIO-ChR2-EYFP. For the presynaptic DN-GH expression experiments, the titers were 1.6 × 10^12^ GC mL^-1^ for AAV9-TRE3G-myr-mScarlet-I, 6 × 10^11^ GC mL^-1^ for AAV9-c-fos-rtTA3G, 4 × 10^12^ GC mL^-1^ for AAV5-CamKIIα-DN-GH-P2A-mEmerald, 6.25 × 10^11^ GC mL^-1^ for AAV9-CamKIIα-iCre, and 1 × 10^12^ GC mL^-1^ for AAV9-EF1α-DIO-ChR2-EYFP. For DN-GH optogenetic engram reactivation experiments, the titers were 7.5 × 10^12^ genome copy (GC) mL^-1^ for AAV9-TRE3G-ChR2-EYFP, 7.5 × 10^12^ GC mL^-1^ for AAV9-c-fos-rtTA3G, and 4 × 10^12^ GC mL^-1^ for AAV9-CamKIIα-DN-GH-P2A-mCherry. For the control experiments, the titers were 7.5 × 10^12^ GC mL^-1^ for AAV9-TRE3G-ChR2-EYFP, 7.5 × 10^12^ GC mL^-1^ for AAV9-c-fos-rtTA3G, and 4 × 10^12^ GC mL^-1^ for AAV9-CamKIIα-mCherry. Due to the potential anterograde transsynaptic expression of AAV serotype 9 at high titer, AAV5 was chosen for the presynaptic DN-GH expression ^41^. The original viruses were diluted with Dulbecco’s phosphate-buffered saline (added NaCl to a final concentration of 10 mM) to match the aforementioned titers.

### Stereotactic surgery

DG viral injections were bilaterally targeted to (AP: −2.1 mm, ML: ± 1.3 mm, DV: −2.0 mm). Optic cannulas (Ferrule material: Black Ceramic, Ferrule diameter: 1.25 mm, Fiber diameter: 300 μm, NA: 0.37, Fiber length: 2 mm, Newdoon) were bilaterally inserted 0.2 mm above the viral injection sites (AP: −2.1 mm, ML: ± 1.3 mm, DV: −1.8 mm). Drug infusion cannulas (C232G-3.0/SPC, 22G, guide length: 2 MM, P1 Technologies) were bilaterally targeted to (AP: −2.1 mm, ML: ± 1.5 mm, DV: −2.0 mm). Medial Entorhinal Cortex (MEC) viral injections were bilaterally targeted to (AP: −4.6 mm, ML: ± 3.5 mm, DV: −3.5 mm). For all experiments other than optogenetic engram reactivation, a 500 nL or 800 nL mixture of virus cocktail was injected into the DG or MEC, respectively. For optogenetic engram reactivation, a 300 nL mixture of virus cocktail was injected into the DG. The stereotaxic surgery was divided into two due to viral titer limits for DN-GH optogenetic engram reactivation and its control experiments.

## Behavior

### Fear conditioning

Animals were conditioned 2-3 weeks after the AAV injection and were single-caged 6 days before the CFC. Each animal was habituated to the hands of the investigator and the anesthesia chamber for 3 minutes each for 5 days before the CFC. On the conditioning day, animals were briefly anesthetized with isoflurane in the anesthesia chamber and 280 μL of 5 mg/mL Doxycycline (in saline) was injected (i.p.) 2 hours before the conditioning. Similarly, following anesthesia with isoflurane, all animals were injected (i.p.) with either anisomycin (Hello Bio) or an identical volume of saline −1, 0, 2, 4 hours after conditioning. For the initial injection, anisomycin was injected with the dosage of 150 mg/kg. The dosage of subsequent 3 injections was lowered to 50 mg/kg ^18^. Anisomycin was freshly dissolved in 1 M HCl and saline and the pH was adjusted to 7.0-7.2 with 1 M NaOH to a final concentration of 10 mg/mL before the experiment. Recombinant human growth hormone (rhGH) (Cell Guidance Systems) was dissolved in distilled water to a final concentration of 0.5 mg/mL. Aliquoted tubes (200 μL) were frozen in the −20 °C freezer until the day of the experiment. The dosage for the injections was 4 mg/kg. The fear conditioning session lasted for 300 s and animals received three 0.75 mA shocks of 2 s duration at 208, 238, and 268 s (Med Associates Inc., St Albans, VT). The baseline freezing level was quantified using the first 180 s of the fear conditioning session. Long-term memory was quantified using the retrieval session that lasted for 180 s without any shocks 2 days after fear conditioning.

### Drug infusion

All animals were conditioned 2 weeks after the cannula implantation. One hour before the CFC paradigm, the animals were briefly anesthetized with isoflurane, and the dummy cannula was replaced with the internal cannula. Next, anisomycin (100 μg/ul) or saline was bilaterally infused (0.8 μl per site, 1.6 μl in total, at the speed of 0.15 μl/min) into the DG. After the infusion, the internal cannula was replaced with the dummy cannula, and the animals were put back into the home cage.

### Optogenetics

All animals were conditioned 2 weeks after the AAV injection and went through identical procedures with the CFC paradigm. On the reactivation day, animals were briefly anesthetized with isoflurane, and the optic fiber was connected to the implanted optic ferrule. The animals recovered from the isoflurane for at least 30 min in their home cages before the optogenetic experiments. After the recovery, ChR2 was stimulated using a 473 nm laser (20 Hz, 20 mW, 15 ms pulse width) for a set period (alternating 3 min OFF and ON sessions, 4 sessions) ^4,42^. Freezing bouts and durations were manually detected by a blinded investigator.

### Perfusion, Cell counting, and Spine density analysis

Animals were anesthetized and perfused with 15 mL of ice-cold phosphate-buffered saline (PBS) and 15 mL of ice-cold 4% PFA (in PBS). The isolated brains of the animals were kept at 4 °C for 24 hours in 4% PFA. After 24 hours, the solution was replaced with a 30% sucrose solution (in PBS) and was kept at 4 °C for 48 hours. Subsequently, the brains were frozen in the −80 °C deep freezer and then cryosectioned for imaging. Engram and non-engram cells in the DG were identified and imaged with a confocal microscope (Stellaris 5 or 8, Leica) equipped with a 20X objective lens. The dendrites in the DG were imaged with a 63X objective lens. We used the Imaris software (Bitplane, Zurich, Switzerland) to process and reconstruct the dendrite filaments. Dendrites expressing mScarlet-I were counted as engrams whereas dendrites exclusively expressing mTurquoise2 were counted as non-engrams. For fluorescent intensity comparison, the mean brightness of the cell body in the granule cell layer was used. All investigators were blinded to the groups throughout the analysis.

## Electrophysiology

### General preparation

Animals were subjected to electrophysiology 36-60 hours after CFC. Animals were anesthetized with intraperitoneal injection of Ketamine/Xylazine mixture. Once deeply anesthetized, transcardial perfusion was performed with 20 mL of ice-cold sucrose cutting solution (210 mM Sucrose, 3 mM KCl, 26 mM NaHCO_3_, 1.25 mM NaH2PO4, 10 mM Glucose, 5 mM MgSO_4_, 0.5 mM CaCl_2_, 3 mM Sodium Ascorbate). Subsequently, hippocampal slices (350 μm) were prepared using a vibratome (VT 1200S; Leica) in the ice-cold sucrose cutting solution and then recovered in 32-34 °C artificial cerebrospinal fluid (ACSF, 2 mM NaCl, 3 mM KCl, 26 mM NaHCO_3_, 1.25 mM NaH_2_PO_4_, 2 mM MgSO_4_, 15 mM Glucose, 2 mM CaCl_2_) for 30 minutes. Next, the chamber containing the slices was transferred to a water tank at room temperature (RT) and recovered for at least 1 hour. All recordings were performed with ACSF at 30-32 °C at a flow rate of 2 mL/min. A 4-6 MΩ tip resistance recording pipette was used for whole-cell patch-clamp recordings. ACSF and the sucrose cutting solution were carbogenated (95% O_2_, 5% CO_2_) before and during usage. Recording data was discarded whenever the resting membrane potential was depolarized above –50 mV or when the access resistance was over 30 MΩ.

## Recording

Capacitance and mEPSC were isolated with 0.5 μM tetrodotoxin (Hello Bio) and 100 μM picrotoxin (Hello Bio) in ACSF and recorded in cells in which the membrane potential was held constant at −70 mV. The frequency of mEPSC was calculated using MiniAnalysis (Synaptosoft Inc., Decatur, GA). The recording pipettes were filled with a potassium gluconate-based internal solution (8 mM NaCl, 130 mM K-Gluconate, 10 mM HEPES, 0.5 mM EGTA, 4 mM MgATP, 0.3 mM Na_3_GTP, 5 mM KCl, 0.1 mM Spermine), and the liquid junction potential was not corrected. The pH of the internal solution was adjusted to 7.2-7.3 with KOH and the osmolarity was set to 285-290 mOsm/L.

PPR and AMPA/NMDA EPSC ratio was acquired with 100 μM picrotoxin (Hello Bio) in ACSF and were recorded in cells in which the membrane potential was held constant at −70 mV. The photostimulation (473 nm, 1 ms width) intensity (−1 mW) was adjusted so that the peak amplitude of AMPAR EPSC was 100-300 pA. NMDAR EPSC was defined as the amplitude of the EPSC 50 ms (40 mV) after the onset of photostimulation. For PPR calculations, 3-5 traces (−70 mV) in 6 s intervals were recorded, and the median PPR was defined as the final PPR value. Inter stimulus interval of 50 and 100 ms was acquired from each cell, and the peak value of the first (P1) and second (P2) EPSC was used to calculate the PPR (P2/P1). For AMPA/NMDA EPSC ratio calculations, 3-5 traces of AMPAR EPSCs (6 s interval, −70 mV), 2-3 traces of evoked EPSCs (6 s interval, 0 mV), 3-5 traces of NMDAR EPSCs (6 s interval, 40 mV), and 3-5 traces of AMPAR EPSCs (6 s interval, −70 mV) were recorded sequentially. The median value was used for the ratio calculations. The recording pipettes were filled with a CeMeSO_3_-based internal solution (8 mM NaCl, 130 mM CeMeSO_3_, 10 mM HEPES, 0.5 mM EGTA, 4 mM MgATP, 0.3 mM Na_3_GTP, 5 mM KCl, 0.1 mM Spermine), and the liquid junction potential of 14 mV was corrected. The pH of the internal solution was adjusted to 7.2-7.3 with KOH and the osmolarity was set to 285-290 mOsm/L.

### Growth hormone internalization

All experiments were performed with 100 μM picrotoxin (Hello Bio) in ACSF. The brain slices (350 μm) were pre-incubated in the ACSF with His-rhGH (100 ng/mL, Sino Biological) for at least 10 minutes before the iTBS (2 s protocol, 8 s delay). The tip of the stimulating electrode was placed in the medial molecular layer, and each stimulation of the iTBS was adjusted so that the peak amplitude of AMPAR EPSC was 400-500 pA. For PPR and sEPSC recordings, after 4 rounds of iTBS the brain slices were further incubated in the rhGH containing ACSF for at least 5 minutes until the sEPSC or PPR recordings started. For His-tag IHC, the brain slices were submerged in ice-cold 4% PFA (in PBS) immediately after 4 rounds of iTBS. Brain slices were fixed in PFA for approximately 16 hours and were subjected to IHC.

### Immunohistochemistry

For c-fos and GH IHC, coronal sections (30 μm) of the frozen brain samples were acquired using a cryostat (Leica). Sections were blocked with a blocking solution containing 5% goat or donkey serum (Rockland) and 0.3% Triton X-100 (Sigma-Aldrich) (in PBS) for 1 hour at RT. For His and GH IHC on the stimulated brain slices (350 μm), samples were blocked in a blocking solution containing 10% goat or donkey serum and 0.5% Triton X-100 (in PBS) for 1 hour at RT. Next, sections were incubated with the appropriate primary antibody at 4 °C overnight [rabbit anti-c-fos 1:1000 (Synaptic Systems, 226 008), goat anti-GH 1:40 (R&D systems, AF1067), and rabbit anti-6XHis tag 1:500 (Abcam, AB9108)]. Following overnight incubation, the sections were incubated with the appropriate secondary antibody for 2 hours at RT and were mounted with Vectashield (Vector Laboratories) [Goat anti-rabbit IgG Alexa Fluor 568 1:500 (Thermo Fisher Scientific, A-11011) and Donkey anti-goat IgG Alexa Fluor 568 1:500 (Thermo Fisher Scientific, A-11057)]

### BONCAT and Western Blot

The procedures of the BONCAT were slightly modified from previous studies ^35-37,43,44^. Prior to the experiment, the animals were put on a methionine-deficient diet (Dooyeol Biotech) for at least a week. The azide-containing non-canonical amino acid L-azidohomoalanine (AHA, Vector Laboratories) 100 μg/gbw was injected (i.p.) 2 hours before the CFC. As previously described, saline or anisomycin were injected with the GH or vehicle −1, 0, and 2 hours following CFC. Four hours after the CFC, mice were deeply anesthetized with isoflurane and sacrificed. The hippocampi were immediately dissected and snap-frozen with liquid nitrogen. The frozen hippocampi were kept in the −80 °C deep freezer until the subsequent experiments. For the BONCAT experiment, hippocampi were homogenized in 1X RIPA buffer (LPS solution) containing a protease inhibitor cocktail (Roche). The protein concentration of the homogenized samples was measured with the BCA assay (Thermo Fisher Scientific). Following the manfacturer’s instructions, the nascent protein synthesis was tagged with the biotin Click iT protein reaction buffer kit (Thermo Fisher Scientific). Briefly, AHA incorporated into the newly synthesized proteins was detected with the biotin alkyne (Vector Laboratories) through a copper-catalyzed click reaction. After the reaction, the samples were resuspended in 1X RIPA with 5X SDS buffer (LPS solution) and were separated using the 4-12% gradient gel. After the separation, the samples were transferred onto a nitrocellulose membrane. The total protein was quantified using the Ponceau staining. The membrane was blocked with 5% skim milk and incubated with HRP-conjugated streptavidin 1:1000 (Thermo Fisher Scientific, N100) at 4 °C overnight. Chemiluminescent signals were imaged and quantified using a ChemiDocTM MP device (Bio-Rad).

### Statistics

All statistical tests were conducted with either custom scripts written in R version 4.2.2 and RStudio version 2022.12.0+353 or GraphPad Prism version 10.2.0 (Graphpad Software, Inc., San Diego, California, USA). The base R *stats* package was used for the Shapiro-Wilk test. *MixOmics* version 6.22.0 was used for the multilevel PCA, ellipses in the score plot indicate the 95% confidence level, multivariate t-distribution. The *ropls* version 1.30.0 was used for the OPLS-DA. Before using parametric tests, all data was tested with the Shapiro-Wilk test. If the data did not follow the normal distribution, the non-parametric counterpart of the tests was used. Post-Hoc tests (Dunn test) were not performed unless the data passed the omnibus test (Kruskal Wallis test). In all cases, two-tailed statistical tests were used. GraphPad Prism, *ggplot2* version 3.4.4, and Adobe Illustrator (Adobe Systems Inc., San Jose, CA, USA) were used to visualize and finalize the figures. All data were analyzed by the experimenter blinded to the experimental conditions. For parametric data sets using parametric statistical tests, the mean ± S.E.M was used for error bars. For non-parametric data sets using non-parametric statistical tests, the median and interquartile range were used. For longitudinal tracking of data (mEPSC, PPR traces, and freezing levels in optogenetic engram reactivation), the mean ± S.E.M is indicated in the line plot.

## Data availability

All data are available from the corresponding author on request.

## Code availability

This paper does not report original code.

## Acknowledgments

This work was supported by the Institute for Basic Science (IBS-R001-D3)

## Author contribution

C.-H.K., H.P., D.H.H., and B.-K.K. contributed to the study design and wrote the paper. C.-H.K. conducted stereotaxic surgery, animal behavior experiments, electrophysiological experiments, functional annotation analysis, and image analysis. H.P. prepared the DNAs, conducted perfusion, immunohistochemistry, western blot, and BONCAT. D.H.H. assisted with electrophysiological experiments. J.C. assisted with functional annotation analysis. N.Y., I.H., Y.J., and J.C. read and commented on the paper.

## Declaration of Interests

The authors declare no competing interests.

## Notes

### Competing Interest Statement

The authors have declared no competing interest.

