## Supplementary Figures for "Growth hormone is required for hippocampal engram cell maturation"

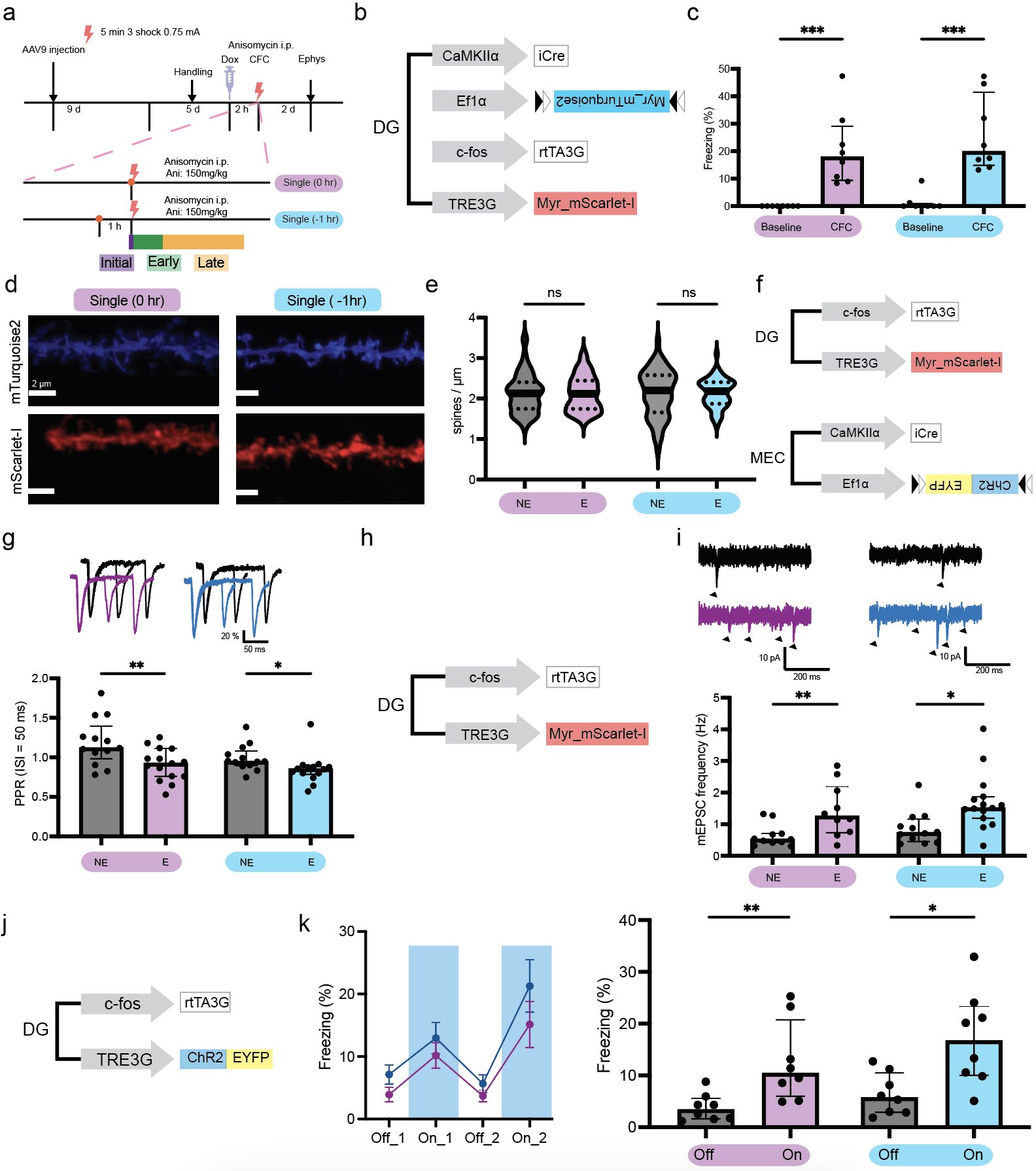

**Extended data Figure. 1: Single anisomycin injection is not sufficient for blocking engram maturation. a**, Experimental procedures for single anisomycin injection. **b**, Viral constructs for natural memory retrieval and spine density analysis. **c**, Behavioral results of the natural memory retrieval (n = 8 animals per group). **d**, Image of the dendrite segments. **e**, Dendritic spine density of neurons (from left to right, n = 61, 62, 62, 60 dendrites). **f**, Viral constructs for PPR analysis. **g**, Top: traces of the PPR. Bottom: PPR (ISI = 50 ms) of neurons (from left to right, n = 13, 15, 13, 14 neurons). **h**, Viral constructs for mEPSC analysis. **i**, Top: traces of the mEPSC. Black arrows indicate each mEPSC spike. Bottom: mEPSC frequency of neurons (from left to right, n = 11, 10, 12, 15 neurons). **j**, Viral constructs for artificial engram reactivation. **k**, Left: behavior paradigm and the quantified freezing behavior. Right: the quantified freezing behavior (n = 8 per group). Mann Whitney U test, ***: P < 0.001, **: P < 0.01, *: P < 0.05.

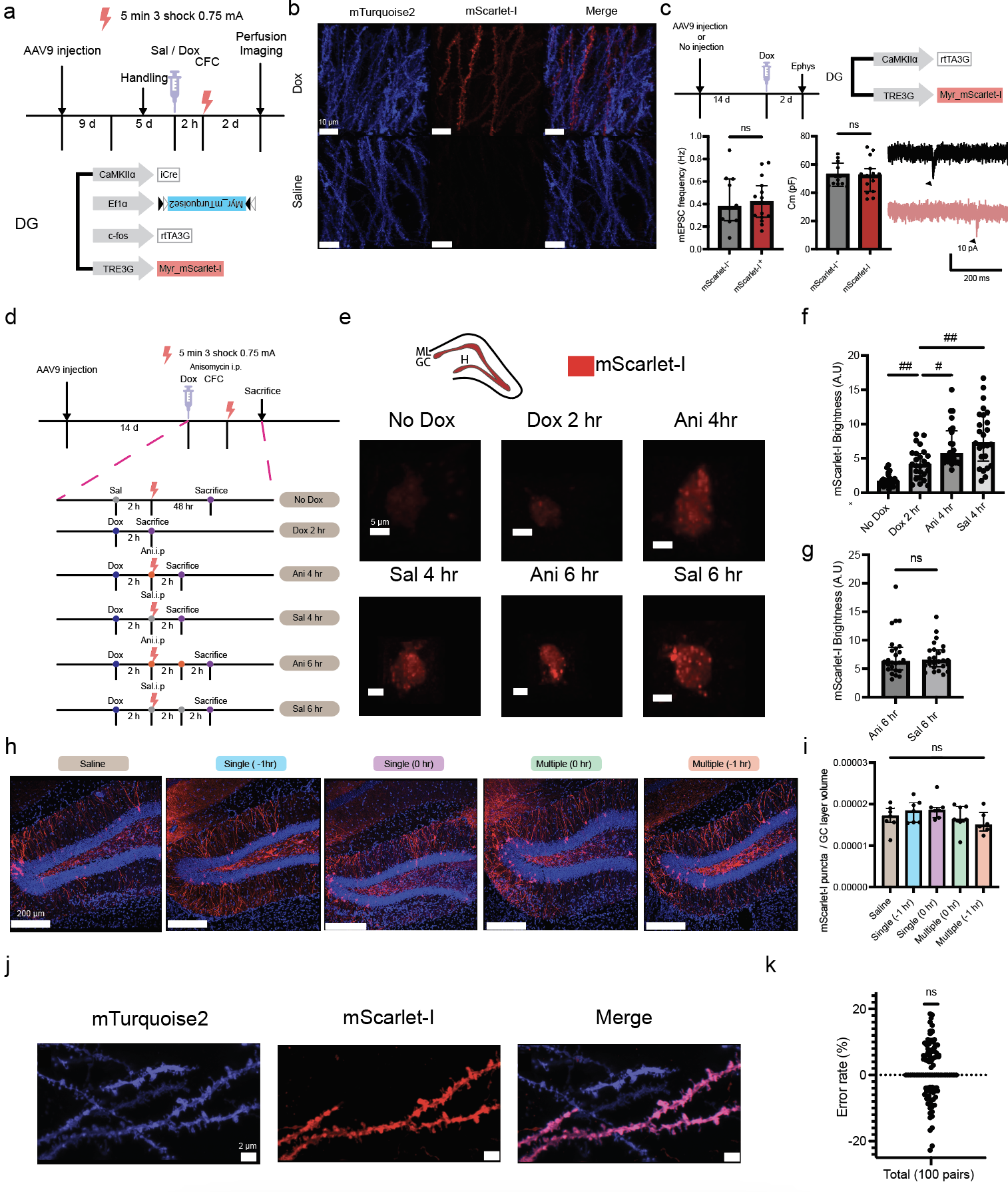

**Extended data Figure. 2: Validation of the engram labeling system.** **a**, Top: experimental procedures used to validate the doxycycline-dependent expression of mScarlet-I. Bottom: viral constructs used for the experiment. **b**, Images of the dendrites in the DG. **c**, Top left: experimental procedures used to test for the effects of viral overexpression on the intrinsic properties of DG cells. Top right: viral constructs used for the experiment. Bottom left**:** mEPSC frequency and the Capacitance of the mScarlet-I positive or negative cells (from left to right, n = 10, 15, 12, 15 neurons). Bottom right: traces of the mEPSC. Black arrows indicate each mEPSC spike. **d**, Experimental procedures for the time course experiment of protein expression. **e**, Images of the cells in the DG. **f-g**, The brightness of the expressed protein (mScarlet-I) in the cell body (n = 25 cells per group). **f**, Dunn test, ##: adj.P < 0.01, #: adj.P < 0.05. **g**, Mann Whitney U test, ns: P > 0.05. **h**, Images of the cells in the DG 2 days after CFC. **i**, The proportion of labeled cells in the DG 2 days after the CFC (from left to right, n = 7, 7, 7, 7, 6 animals). Kruskal Wallis test, ns: P > 0.05. **j**, Representative images of the dendrites. **k**, The error rate of spine counting (Error rate: (mScarlet-I puncta – mTurquoise2 puncta) / mTurquoise2 puncta). Shapiro Wilk test P > 0.05, One sample t-test (hypothetical mean = 0), ns: P > 0.05.

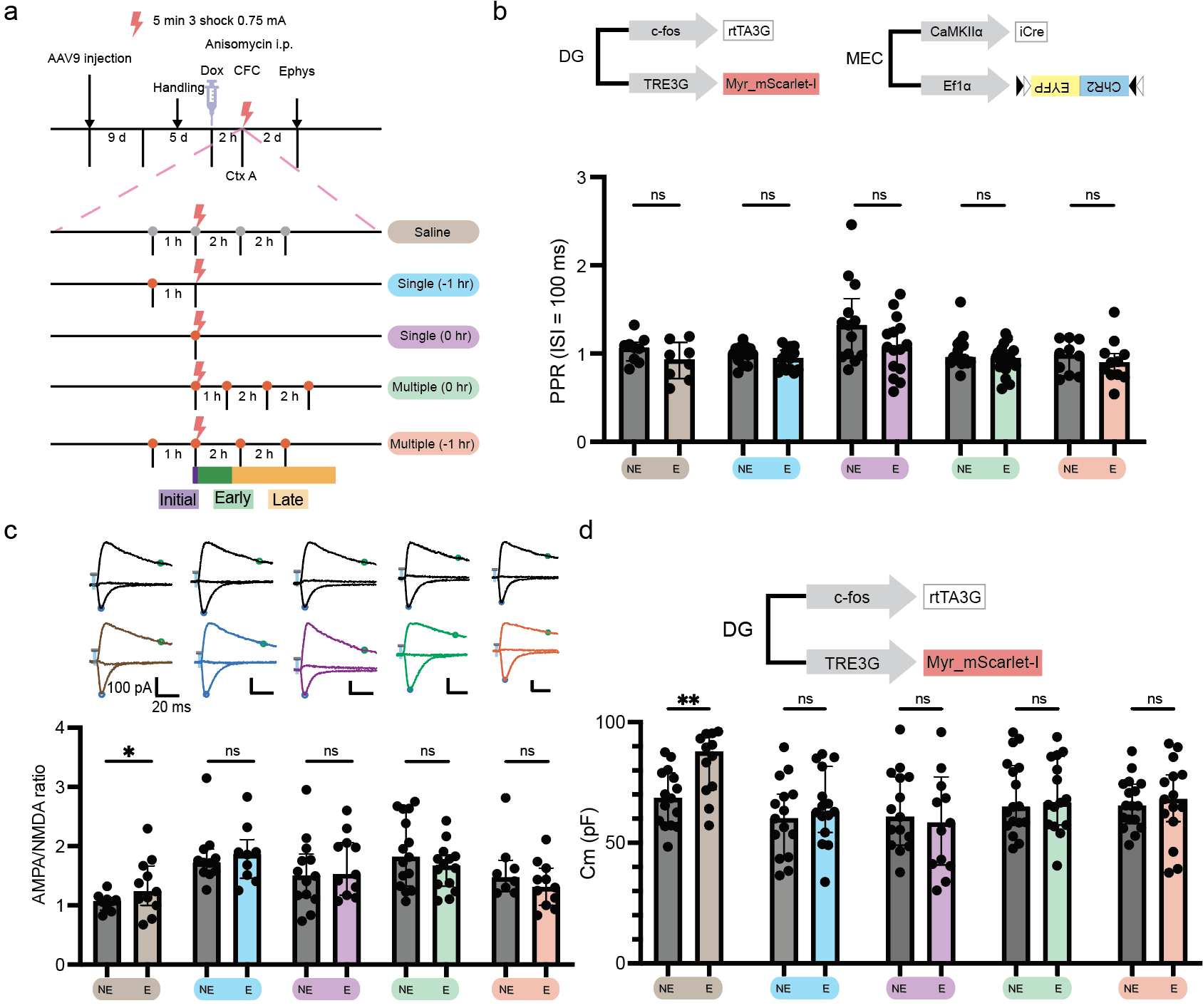

**Extended data Figure. 3: Initial translation-independent electrophysiological properties.** **a**, Experimental procedures for the behavioral paradigm used to block engram maturation. **b**, Top: viral constructs used to measure evoked properties (AMPA/NMDA ratio and PPR). Bottom: PPR (ISI = 100 ms) of neurons (from left to right, n = 9, 8, 13, 14, 13, 15, 16, 18, 10, 11 neurons). **c**, Top: AMPA/NMDA ratio traces acquired at +40 mV, 0 mV, and -70 mV. Bottom: AMPA/NMDA ratio of neurons (from left to right, n = 11, 9, 11, 9, 13, 11, 15, 14, 8, 11 neurons). **d**, Top: viral constructs used for capacitance measurement. Bottom: capacitance of neurons (from left to right, n = 15, 12, 15, 15, 15, 13, 17, 15, 17, 15 neurons). Mann Whitney U test, **: P < 0.01, *: P < 0.05, ns: P > 0.05.

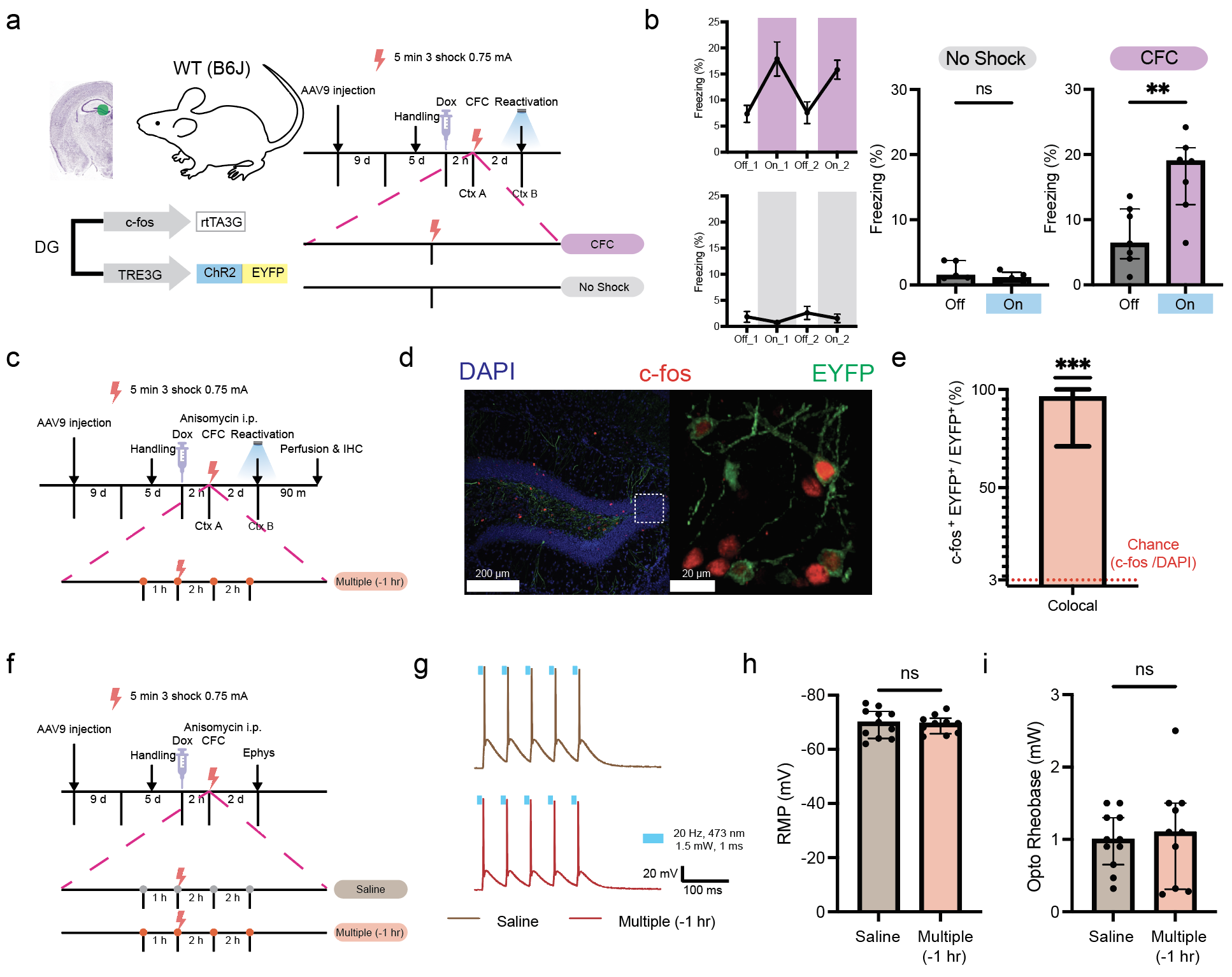

**Extended data Figure. 4: Validation of the optogenetic engram reactivation. a**, Experimental procedures to test the aversive memory dependence of engram reactivation. **b**, Left: behavior paradigm and the quantified freezing behavior. Right: the quantified freezing behavior (no shock: n = 5, CFC: n = 7 animals). **c**, Experimental procedures for the behavioral paradigm used for colocalization analysis of c-fos and ChR2-EYFP. **d**, Images of the DG after c-fos IHC. **e**, Colocalization percentage of c-fos and ChR2-EYFP (n = 12 brain slices from 3 animals). The red dotted line indicates the median of the chance level (c-fos/DAPI). **f**, Experimental procedures for the behavioral paradigm used to test the functionality of ChR2-EYFP. **g**, Traces of the ChR2-EYFP mediated 20 Hz stimulation. **h**, The resting membrane potential (RMP) of the neurons (from left to right, n = 11, 10). **i**, Minimum laser intensity to elicit an action potential (Opto-Rheobase) (from left to right, n = 11, 10). Mann Whitney U test, ***: P < 0.001, **: P < 0.01, ns: P > 0.05.

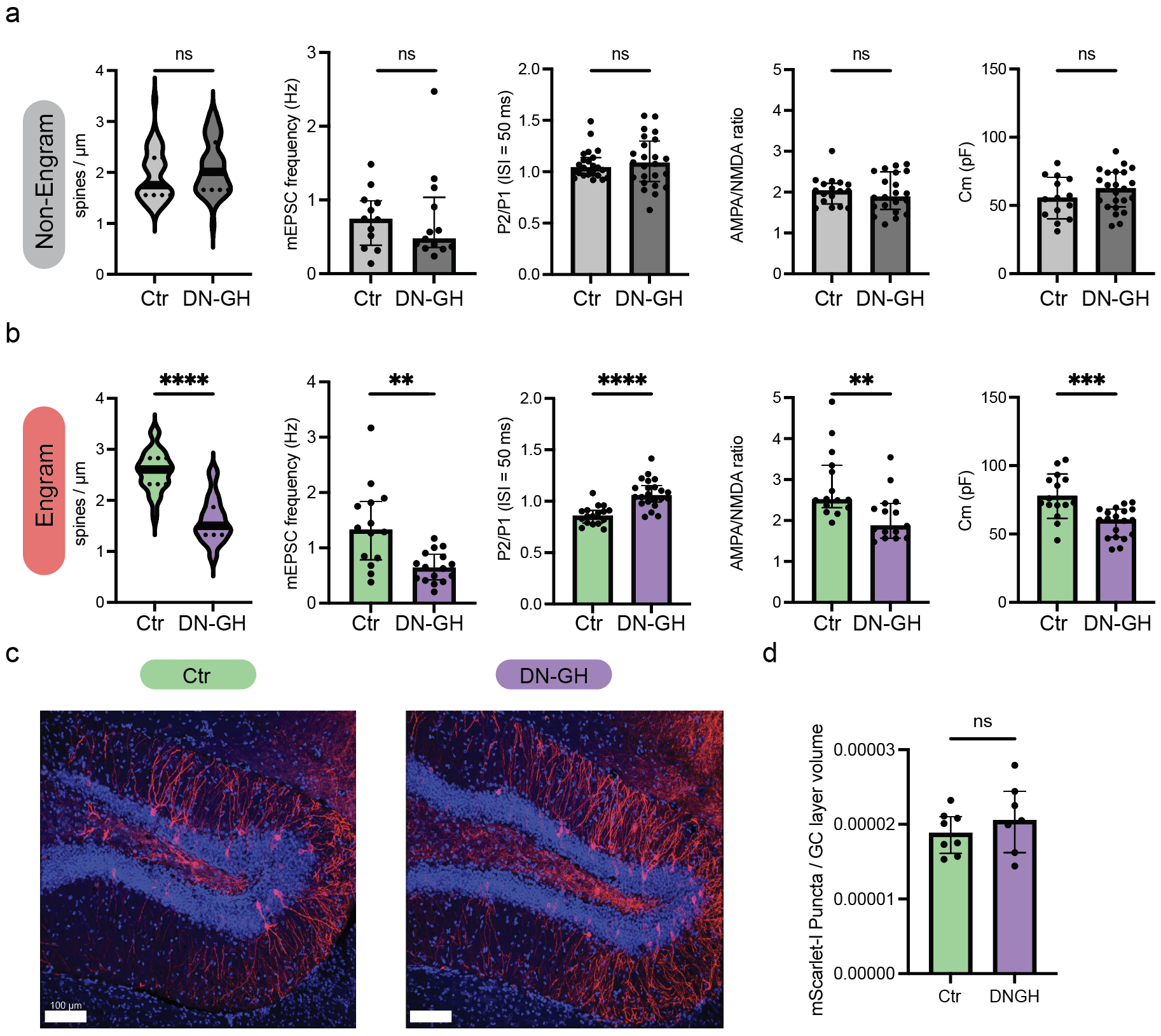

**Extended data Figure. 5: The specificity of the DN-GH on engram cells. a**, Comparison of non-engram neurons or dendrites (from left to right, n = 41, 51, 12, 13, 23, 24, 17, 21, 14, 23). **b**, Comparison of engram neurons or dendrites (from left to right, n = 41, 50, 14, 16, 17, 22, 15, 15, 15, 19). **c**, Representative image of the DAPI stained DG expressing mScarlet-I. **d**, The proportion of mScarlet-I labeled cells in the DG (Ctr: n= 8, DN-GH: n = 7). Mann Whitney U test, ****: P < 0.0001, ***: P < 0.001, **: P < 0.01, ns: P > 0.05.

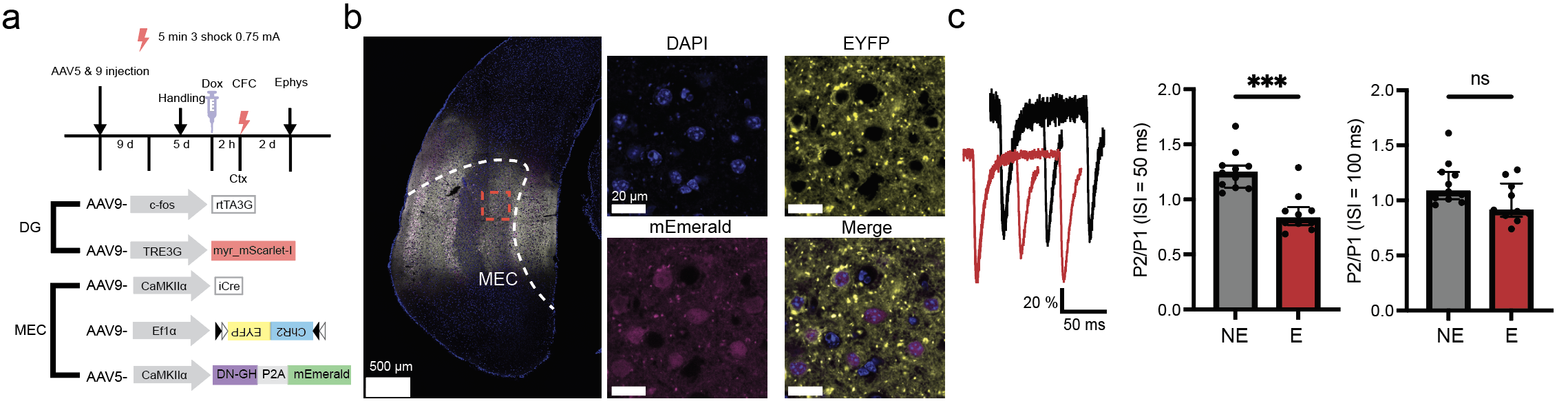

**Extended data Figure. 6: The effect of DN-GH expression in the MEC. a**. Top: Experimental procedures for the presynaptic DN-GH expression and ex vivo electrophysiology. Bottom: viral constructs for presynaptic DN-GH expression. **b**, Images of the MEC. **c**, Left: traces of the PPR. Right: PPR (ISI = 50 or 100 ms) of neurons (from left to right, n = 11, 10, 11, 10 neurons). Mann Whitney U test, ***: P < 0.001, ns: P > 0.05.

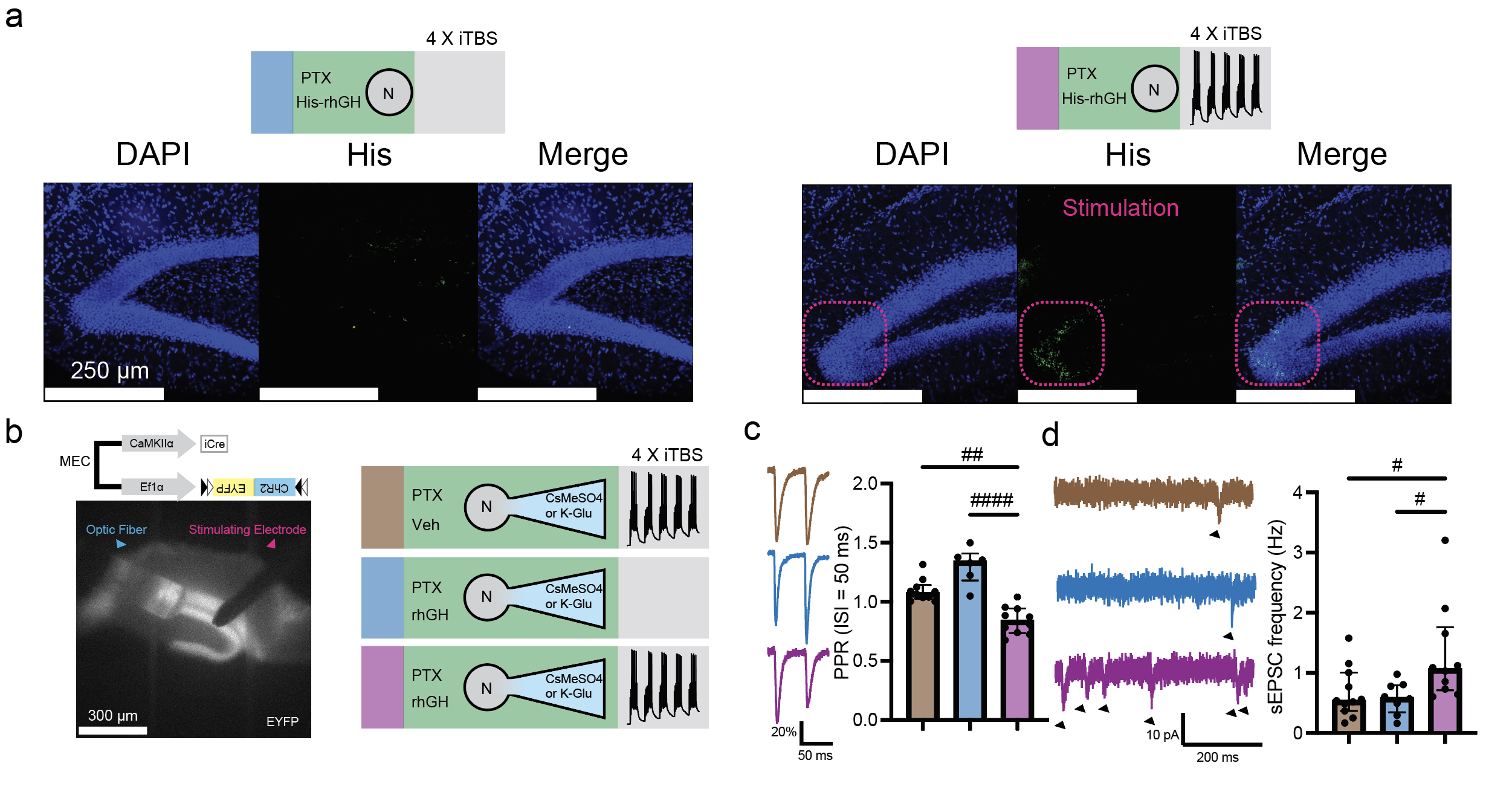

**Extended data Figure. 7: Activity-dependent internalization of external GH. a**, Image of the DG after his-tag IHC. **b**, schematic for PPR and sEPSC analysis after GH internalization. **c**, Left: traces of the PPR. Right: comparison of the PPR (from left to right, n = 10, 6, 10 neurons). **d**, Left: traces of the sEPSC. Right: comparison of the sEPSC frequency (from left to right, n = 12, 8, 10 neurons). Dunn test, ####: adj.P < 0.0001, ##: adj.P < 0.01, #: adj.P < 0.05, ns: adj.P > 0.05.

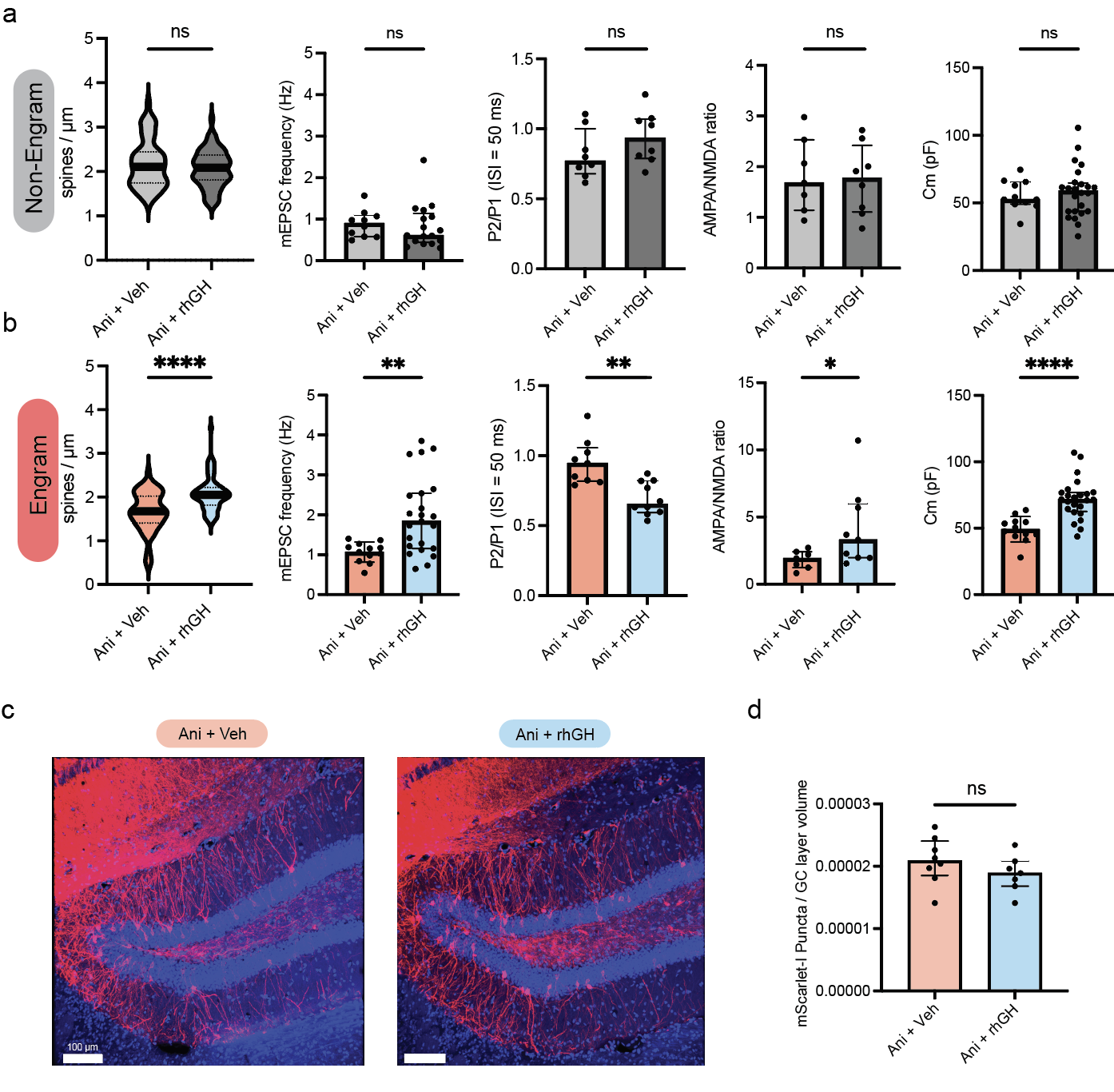

**Extended data Figure. 8: The specificity of the rhGH on engram cells. a**, Comparison of non-engram neurons or dendrites (from left to right, n = 61, 67, 12, 13, 23, 24, 17, 21, 14, 23). **b**, Comparison of engram neurons or dendrites (from left to right, n = 60, 71, 14, 16, 17, 22, 15, 15, 15, 19). **c**, Image of the DAPI stained DG expressing mScarlet-I. **d**, The proportion of mScarlet-I labeled cells in the DG (Ani+Veh: n= 8, Ani+rhGH: n = 7). Mann Whitney U test, ****: P < 0.0001, **: P < 0.01, ns: P > 0.05.
